# Running Promotes Spatial Bias Independently of Adult Neurogenesis

**DOI:** 10.1101/125260

**Authors:** Jason S. Snyder, Shaina P. Cahill, Paul W. Frankland

## Abstract

Different memory systems offer distinct advantages to navigational behavior. The hippocampus forms complex associations between environmental stimuli, enabling flexible navigation through space. In contrast, the dorsal striatum associates discrete cues and favorable behavioral responses, enabling habit-like, automated navigation. While these two systems often complement one another, there are instances where striatal-dependent responses (e.g. approach a cue) conflict with hippocampal representations of spatial goals. In conflict situations, preference for spatial vs. response strategies varies across individuals and depends on previous experience, plasticity and the integrity of these two memory systems. Here, we investigated the role of adult hippocampal neurogenesis and exercise on mouse search strategies in a water maze task that can be solved with either a hippocampal-dependent place strategy or a striatal-dependent cue-response strategy. We predicted that inhibiting adult neurogenesis would impair hippocampal function and shift behavior towards striatal-dependent cue responses. However, blocking neurogenesis in a transgenic nestin-TK mouse did not affect strategy choice. We then investigated whether a pro-neurogenic stimulus, running, would bias mice towards hippocampal-dependent spatial strategies. While running indeed promoted spatial strategies, it did so even when neurogenesis was inhibited in nestin-TK mice. These findings indicate that exercise-induced increases in neurogenesis are not always required for enhanced cognitive function. Furthermore, our data identify exercise as a potentially useful strategy for promoting flexible, cognitive forms of memory in habit-related disorders that are characterized by excessive responding to discrete cues.

## INTRODUCTION

Multiple memory systems theory holds that different brain regions support distinct types of learning. The extent to which a given memory system exerts control over behavior depends on the types of sensory stimuli present and the extent to which they can make predictions about the future (White and McDonald, 2002). Prior experience, emotion and hormones are some of the many factors that can also influence which memory system, and associated strategies, are employed to solve a given problem. Two systems that mediate distinct and complementary types of learning are the hippocampal and dorsal striatal memory systems. The hippocampus forms a rich set of associations between sensory stimuli of all modalities (often referred to as cognitive memory), enabling flexibility in goal-directed behavior (Schiller et al., 2015). For example, by forming associations between multiple different cues in an environment, the hippocampus enables rodents to navigate to a specific place in an environment (Morris et al., 1982; Eichenbaum et al., 1990). In contrast, the dorsal striatum supports stimulus-response behaviors, where a subject learns to perform a specific response to a discrete cue (often referred to as habit memory)(Packard et al., 1989). Examples include learning to approach a specific cue, or make a specific body turn at a choice point, in order to receive a reward. While stimulus-response learning is often referred to as dorsal striatum-dependent, it is primarily a function of the *dorsolateral* striatum (Devan and White, 1999) (but see (Ferbinteanu, 2016).

Many of the factors that regulate the relative use of these two memory systems have been identified using “dual solution” navigational tasks in rodents, which can be solved through either spatial or response strategies. When these strategies are congruent, dysfunction in either the hippocampus or dorsal striatum does not affect performance since the animal can rely on the other system to solve the task. However, the interactive nature of these two memory systems can be revealed when, after a period of training, task modifications cause spatial and response strategies to produce conflicting outcomes. In this scenario, the subject must choose between strategies. Compromising one memory system with lesions, inactivation or NMDA receptor blockade promotes use of the alternate strategy (i.e. hippocampal dysfunction increases striatal-based cue-responding and vice versa; (McDonald and White, 1994; Packard and McGaugh, 1996; Packard and Teather, 1997). Conversely, dopamine agonists promote spatial strategies when injected into the hippocampus and they promote response strategies when injected into the dorsal striatum (Packard and White, 1991).

In dual solution paradigms, stimulus-response strategies are typically adopted with additional training; i.e. performance becomes automatic and habit-like (Packard and McGaugh, 1996). Stress (Kim et al., 2001; Schwabe et al., 2010) and anxiogenic drugs (Wingard and Packard, 2008) also promote response strategies and rats with high trait anxiety are more likely to display cued, instead of place, strategies (Hawley et al., 2011). In contrast, estrogen promotes hippocampal-dependent place strategies in dual solution paradigms (Korol and Pisani, 2015). However other factors that specifically promote spatial/hippocampal strategies in dual solution tasks are less well known. Moreover, little is known about the neural cell types that are responsible for modulating hippocampal-dorsal striatal bias.

Exercise is one factor that has profound effects on hippocampal structure and function. Exercise is associated with increased hippocampal volume, cellular plasticity, molecular signalling changes and memory improvements in both humans and rodents (Voss et al., 2013; Hamilton and Rhodes, 2015). Moreover, while exercise is associated with increased dorsal striatal volume in adolescent humans (Chaddock et al., 2010) its effects in adults are limited to the hippocampus (Erickson et al., 2011). This may suggest greater effects on hippocampal-dependent behaviors however, in rats, exercise enhances both place and response learning in single solution learning tasks (Korol et al., 2013). Whether it promotes one strategy over another in a dual solution paradigm has not been tested.

While exercise has diverse effects, it has a unique impact on hippocampal circuits by increasing the proliferation, maturation and survival of new dentate gyrus neurons (van Praag et al., 1999b; van Praag, 2008; Snyder et al., 2009b; Piatti et al., 2011). Adult-born neurons have enhanced plasticity and may play a powerful role in hippocampal behavior (Ge et al., 2007; Tronel et al., 2010; Snyder and Cameron, 2012). Since spatially-trained rats have greater neuronal survival than cue-trained rats (Gould et al., 1999; Epp et al., 2007), and strategy choice correlates with measures of new neuron activation (Yagi et al., 2015), neurogenesis functions may be apparent in dual solution paradigms. We therefore first tested the hypothesis that adult-born neurons promote hippocampal-dependent spatial strategies in a dual solution task. Contrary to our prediction, blocking neurogenesis in a transgenic mouse model did not affect place vs cue choices in a water maze paradigm. Second, we tested the hypothesis that spatial bias is most pronounced when neurogenesis is enhanced following exercise. Consistent with our prediction, running promoted hippocampal-dependent spatial search strategies. However, running-induced spatial bias persisted even when adult neurogenesis was suppressed. These findings may be relevant for understanding how cognitive strategies can be harnessed to offset striatal-dependent habit disorders. They also suggest that, in the absence of direct tests, caution is warranted when interpreting exercise effects as neurogenesis-dependent.

## METHODS

### Animals and treatments

All experiments were approved by the animal care committee at the Hospital for Sick Children and conducted in accordance with the Canadian Council on Animal Care. Mice were housed 3-5 per cage with ad libitum access to food and water. All testing was performed during the light phase (12 hr light-dark cycle with lights on at 7:00 am). Male wild type mice were generated by crossing C57Bl/6NTac and 129S6/SvEv strains (Taconic). To suppress adult neurogenesis we used a transgenic line that expresses herpes simplex virus thymidine kinase (HSV-TK) under the control of the nestin promoter (TK mice). In TK models, viral thymidine kinase phosphorylates the antiviral drug ganciclovir, a nucleotide analog, which then interferes with DNA synthesis and causes death of dividing TK-expressing cells. By expressing TK in hippocampal precursor cells neurogenesis can be selectively inhibited in adulthood (Garcia et al., 2004; Saxe et al., 2006; Deng et al., 2009; Singer et al., 2011; Snyder et al., 2011; Cummings et al., 2014; Snyder et al., 2016). TK mice were backcrossed onto a C57Bl/6 background and crossed with 129S6/SvEv mice (Taconic) to obtain male and female experimental mice (both sexes distributed equally across experimental groups). Estrous cycle was not monitored in the female mice. Wild type (WT) and TK mice were given the antiviral prodrug valganciclovir ad libitum in powdered food (0.08%, ∼70 mg/kg daily) from 6-12 weeks of age, at which point they underwent behavioral testing. To enhance neurogenesis, in some experiments mice were given access to running wheels (Med Associates) for 4 weeks prior to behavioral testing. Some mice were given the thymidine analog bromodeoxyuridine (BrdU) to label adult-born neurons (200 mg/kg, intraperitoneal, dissolved 10 mg/ml in saline).

### Behavioral testing

We used a dual solution water maze paradigm to investigate place vs. cue navigational preferences (McDonald and White, 1994; Kim et al., 2001). Testing was performed in a circular pool (120 cm diameter) that was filled to 30 cm from the top of the pool with 23°C water. A white curtain, 1.5 m from the pool edge, surrounded the pool. Four simple shapes, 50-100 cm in width and approximately 100-150 cm above the pool surface, were present at various places on the curtain and served to provide the mice with distal cues for solving the task. Importantly, the position of the curtain and distal cues remained fixed throughout the entire study. Pool water was made opaque with white paint and during the training phase a circular escape platform (10 cm wide, submerged 1 cm below the water surface) was placed in the center of the NW quadrant and a cue hung directly above the platform, thereby reliably predicting its location (15 ml cylindrical Falcon tube with black and white stripes; Fig. 1A). Mice received 4 trials/day (∼5 min ITI) and were released from the N,S,E, or W points of the pool in a pseudorandom fashion. If a mouse did not escape within 60 sec it was placed on the platform for 10 sec before being returned to its cage until the next trial. Groups of mice were trained for 3, 6, or 9 days to develop the task parameters, after which 9 days of training was used for all subsequent experiments. The testing order of different experimental groups was counterbalanced across the day. Performance was measured as the latency to reach the platform.

**Figure 1:**
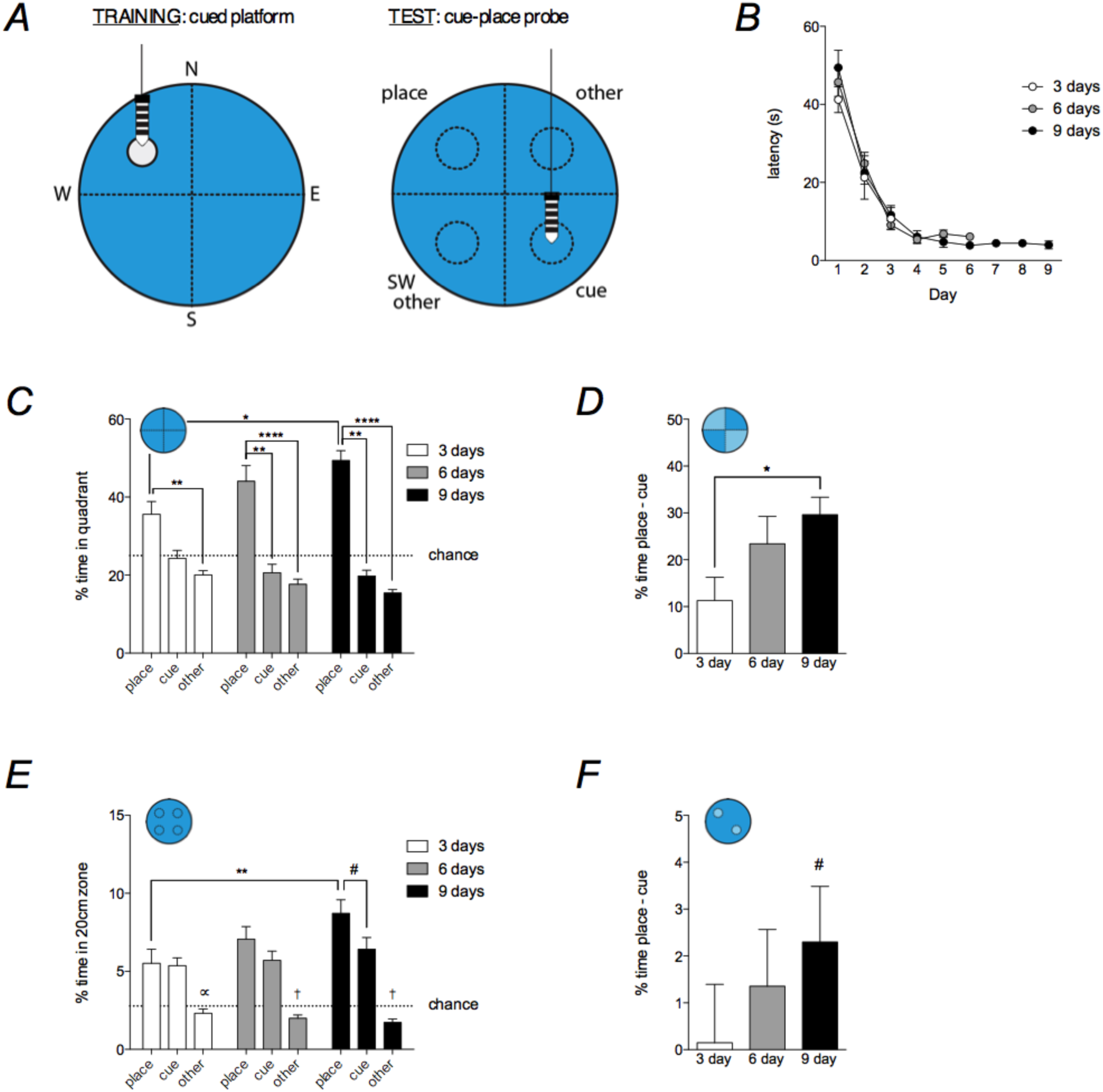
Dual solution water maze paradigm. <u>**A**</u>) Training: mice learn to swim to a cued escape platform. Test: the platform is removed and the cue is moved to the opposite quadrant. Mice are free to choose between the former platform place and the new cue location. Compass locations indicate release points. Dashed lines indicate quadrants and 20 cm zones. <u>**B**</u>) Mice escaped to the platform faster on each day during the first 3 days of training with no differences between groups (effect of day: F_2,52_ = 127, P < 0.0001, effect of group: F_2,26_ = 0.7, P = 0.7; differences between days 1, 2, 3 all P < 0.0001). Latencies did not improve after day 3 in the 6 and 9-day groups (days 1-6, effect of day: F_5,95_ = 134, P < 0.0001; differences between days 4-6 all Ps > 0.1). <u>**C**</u>) On cue-place probe trials, mice spent more time searching in the place quadrant than the cued and “other” quadrants (within group Kruskal Wallis tests all P < 0.01; **P < 0.01, ****P < 0.0001). Time spent searching in the place quadrant was greater in the 9 day group compared to the the 3 day group (ANOVA F_2,26_ = 4.2, *P < 0.05) <u>**D**</u>) Spatial bias in the cue-place probe test expressed as a difference score. Mice showed progressively greater bias for the spatial location with additional days of training. Nine days of training significantly increased spatial bias relative to 3 days of training (ANOVA F_2,26_ = 3.4, P < 0.05, *P < 0.05). The 6 and 9 day groups searched in the spatial location significantly more than chance (one sample t-tests: 6 days T_9_ = 4.0, P < 0.05, 9 days T_10_ = 7.9, P < 0.0001; 95% confidence intervals, 6 days: 10.2 – 36.6, 9 days: 21.2 – 38.0). <u>**E**</u>) Probe trial search patterns in 20 cm zones. Mice spent more time searching in the place and cue zones than in the “other” zones (effect of zone F_2,52_ = 43, P < 0.0001; effect of training duration F_2,26_ = 4, P = 0.03; interaction F_4,52_ = 2, P = 0.17; comparison between “other” zones and place/cue: ∝P < 0.05, ^†^P < 0.001). Time spent in the place and cue zones was greater than chance (one sample t-tests, all Ps < 0.001). Time spent in the place zone was greater in the 9 day group than in the 3 day group (**P < 0.01). There was a trend for more time spent in the place zone than in the cued zone in the 9 day group (^#^P = 0.07). <u>**F**</u>) Additional days of training did not significantly increase the spatial bias difference score (ANOVA F_2,26_ = 0.7, P = 0.52), though 9 days of training resulted in a tendency to spend more than chance amounts of time in the place zone (one sample t-test T_8_ = 1.9, ^#^P = 0.09; 95% confidence interval: -0.4 – 5.0). N = 8-12/group.

To assess strategy choice, all groups were given a 60 sec cue-place probe trial one day after the training phase. For the probe trials the platform was removed, the cue was moved to the opposite quadrant (SE) and mice were released from the SW point of the pool. The testing environment was otherwise unaltered relative to the training phase. The time spent searching in the NW quadrant that formerly contained the platform typically reflects hippocampal-dependent, cognitive, place memory and the time spent searching in the SE quadrant that contained the cue reflects striatum-dependent, stimulus-response, habit memory (McDonald and White, 1994). The time spent searching the NE and SW quadrants, which never contained the platform or the cue, was averaged and collectively referred to as the “other” quadrant. In initial experiments we also examined the amount of time spent searching in smaller, 20 cm diameter circular zones that centered on the former platform location, current cue location, or equivalent locations in the SW and NE areas of the pool (Fig. 1A). In addition to individual quadrant/zone analyses, we also expressed spatial bias as a difference score: the percent time spent in the place quadrant/zone minus the percent time spent in the cued quadrant/zone. Here, values above 0 reflect more time spent in the spatial location over the cued location.

### Histology and assessment of adult neurogenesis levels

Following behavioral testing mice were perfused with 4% paraformaldehyde, brains were post-fixed for an additional 48 hours and then cut coronally at 40 *μ*m on a vibratome throughout the entire length of the hippocampus. Neurogenesis was assessed by immunostaining for the microtubule-associated protein doublecortin (DCX) or the thymidine analog bromodeoxyuridine (BrdU). DCX immunostaining was performed on free-floating sections with fluorescence detection. Briefly, sections were incubated in 0.1 M PBS containing 0.3% triton-x, 3% normal donkey serum, and goat anti-doublecortin (Santa Cruz, sc-8066) primary antibody at 1:200 dilution for 3 days. Sections were washed with PBS and then incubated with Alexa488-conjugated donkey anti-goat secondary antibody (Thermofisher; 1:250 in PBS) for 2 hours, counterstained with DAPI and coverslipped with PVA-DABCO. Doublecortin levels were qualitatively assessed from 2-3 sections, to ensure efficacy of the Nestin-TK mouse model (doublecortin was nearly completely absent from the Nestin-TK mice). BrdU immunostaining was performed on a 1 in 6 series of sections throughout the full extent of the hippocampus. Sections were mounted onto slides, heated to 90°C in citric acid (0.1 M, pH 6.0), permeabilized with trypsin, incubated in 2 N HCl for 30 min to denature DNA, and then incubated overnight with mouse anti-BrdU primary antibody (BD Biosciences, 347580; 1:200 in 0.3% triton-x and 3% horse serum). Sections were then washed and incubated with biotinylated goat anti-mouse secondary antibody (Sigma, B0529; 1:250) for 1 hour and cells were visualized with an avidin-biotin-HRP kit (Vector Laboratories) and cobalt-DAB detection (Sigma Fast tablets). All BrdU-positive cells located in the granule cell layer and adjacent subgranular zone (bilaterally) were counted using a 40x objective, using stereological principles. Counts were multiplied by 6 to estimate the total number of labelled adult-born neurons in WT and TK mice that ran or remained sedentary in the final experiment shown in Fig.4.

### Statistical Analyses

Group differences in water maze performance and neurogenesis levels were assessed by ANOVA (repeated measures where appropriate), followed by Holm-Sidak multiple comparison testing. Differences in quadrant search patterns in the cue-place probe tests were detected by a nonparametric Kruskal-Wallis test, followed by Dunn’s multiple comparison testing. Group differences in spatial bias, measured by the quadrant difference score, were assessed by unpaired t-test. In all cases significance was set at P < 0.05, but statistical trends are noted (0.10 > P < 0.05). All graphs report group means and standard error.

## RESULTS

### Effects of training duration on search strategy

Mice were trained for 3, 6 or 9 days on the cued water maze (N = 8-12/group). Latency to reach the platform decreased over the first 3 days of training and plateaued at approximately 5 sec thereafter (Fig. 1B). There were no group differences in acquisition latency and swim speeds were similar across groups (day 1-3 mean: 3 day: 20.3 cm/s, 6 day: 20.6 cm/s, 9 day: 21.6 cm/s; ANOVA, group effect F_2,26_ = 0.98, P = 0.4). On the probe trial we found that additional days of training promoted a spatial search bias in the quadrant analyses: compared to mice that were trained for 3 days, the 9 day group spent more time searching in the place quadrant (Fig. 1C). Mice spent chance, or lower, amounts of time searching in the cue and “other” quadrants and the time spent in these quadrants tended to decrease with days of training, though decreases were not statistically significant. The spatial bias difference score (% additional time searching in the spatial quadrant than the cued quadrant) increased from 11% to 23% and 30% after 3, 6 and 9 days of training, respectively (Fig. 1D). The spatial bias difference score was greater than zero in both the 6 and 9 day groups, and spatial bias was increased after 9 days of training compared to 3 days of training.

Quadrant-level analyses clearly captured spatial search but cued quadrant search times were lower. We reasoned that smaller 20 cm zones may reveal directed searches near a proximal cue. Indeed, search time in the cued 20 cm zone was above chance and greater than the “other” zone. After 9 days of training mice spent more time in the 20 cm place zone than the 3-day group, and there was a non-significant trend for a spatial bias (place vs cue zones, zone difference score; Fig. 1E-F). Thus, mice used both spatial and cued search strategies but the spatial strategy predominated, particularly after 9 days of training.

We performed quadrant-level analyses for subsequent experiments since this best captured spatial performance and changes in search strategy. We note that, while cued quadrant search time appears low and is sometimes comparable to “other” quadrant search, this actually reflects substantial search in the cued location since, in the absence of a cue, mice spent significantly *less* time in the cued quadrant than in the “other” quadrant (see below, Fig. 3E). Our quadrant analyses are therefore sensitive to both place and cue search patterns.

**Figure 2:**
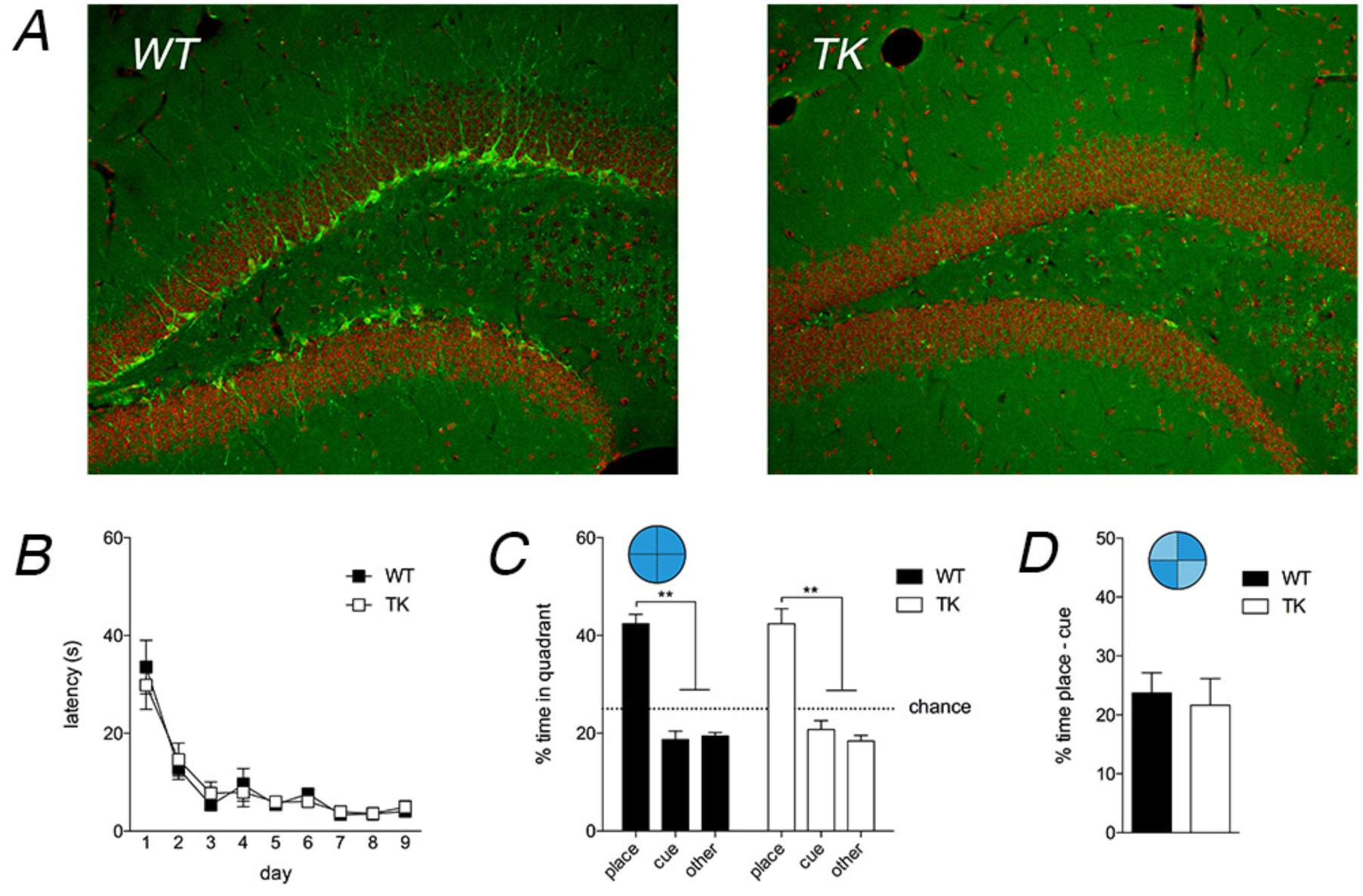
Neurogenesis-deficient TK mice marker doublecortin revealed substantial adult neurogenesis in WT mice but near complete loss of adult-born neurons in TK mice. <u>**A**</u>) Immunostaining for the immature neuronal marker doublecortin revealed adult-born neurons along the deep aspect of the granule cell layer in WT mice (green cell bodies and dendrites) but not in neurogenesis-deficient TK mice. <u>**B**</u>) Acquisition in the cued water maze was similar in WT and TK mice (effect of genotype F_1,146_ = 0.001, P = 0.97). <u>**C**</u>) In probe trial quadrant search, WT and TK mice did not differ and both genotypes spent more time searching in the place quadrant than in the cued or “other” quadrants (Kruskal Wallis tests P < 0.0001, **P < 0.01). <u>**D**</u>) WT and TK mice displayed similar spatial bias as measured by the quadrant difference score (T_19_ = 0.2, P = 0.8). N = 10-11/group.

**Figure 3:**
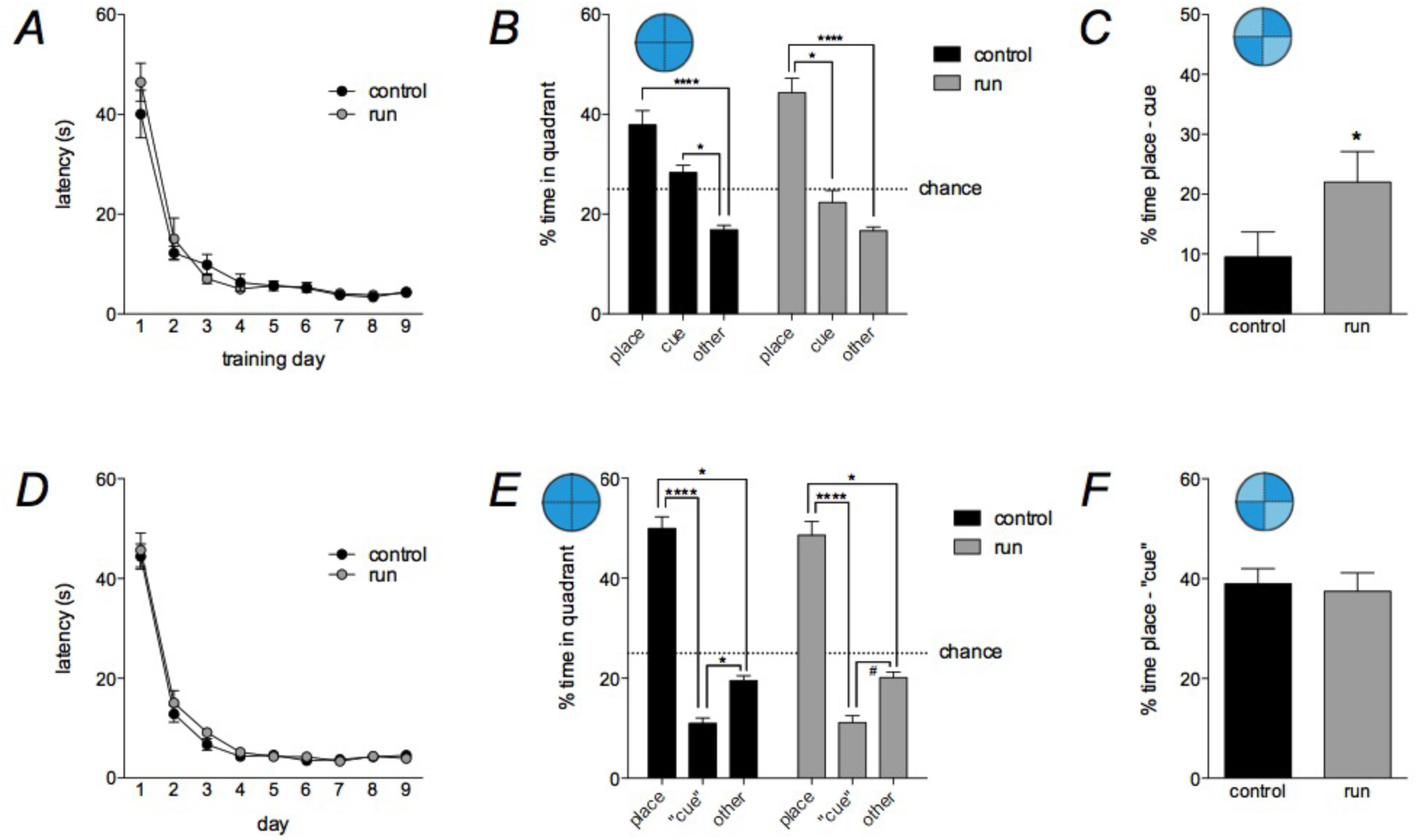
Four weeks of running promotes spatial bias in the cue-place probe test. <u>**A**</u>) Running did not alter escape latencies during cued training (effect of running F_1,15_ = 0.4, P = 0.6). <u>**B**</u>) In the probe trial, all mice spent significantly more time searching in the place quadrant than the “other” quadrants. However, control mice also searched more in the cued quadrant than the “other” quadrant. Conversely, running mice spent more time in the place quadrant than in the cued quadrant (Kruskal Wallis tests P < 0.001, *P < 0.05, ****P < 0.0001). <u>**C**</u>) As measured by the quadrant difference score, running increased spatial bias in the cue-place probe test (T_15_ = 1.9, *P < 0.05). N = 8-9/group. <u>**D**</u>) Running did not alter cued water maze acquisition in a second cohort of mice that were subsequently tested in an uncued, place-only probe trial (effect of running F_1,20_ = 0.6, P = 0.4). <u>**E**</u>) In the place-only (uncued) probe trial all mice preferred the place quadrant and avoided the opposite, “cued” quadrant (i.e. where the cue was located in all other experiment probe trials; Kruskal Wallis tests *P < 0.05, ****P < 0.0001, ^#^P = 0.08). <u>**F**</u>) Running mice were not different from controls in terms of their spatial preference as measured by a quadrant difference score – both groups showed strong spatial memory. The graph shows time in NW minus time in SE, even though a cue was not present in SE, for consistency with the other datasets (T_20_ = 0.9, P = 0.4). N = 10-12/group.

### Inhibiting adult neurogenesis does not alter strategy preference

We hypothesized that inhibiting adult neurogenesis would compromise hippocampal function and reduce spatial search strategies when mice are given a choice between place- and cue-dependent search strategies. We therefore chose the 9-day training paradigm since this produced a robust spatial bias that could potentially be reduced on the probe trial. WT and TK mice were treated with valganciclovir for 6 weeks to block adult neurogenesis (Fig. 2A). Mice were then trained on the cued water maze (N = 10-11/group). Acquisition latencies were similar in both WT and TK mice (Fig. 2B) as were swim speeds (WT 19.0 cm/s, TK 19.1 cm/s; ANOVA effect of genotype F_1,18_ = 0.14, P = 0.7). Both genotypes displayed similarly high levels of spatial quadrant search on the probe trial. Moreover, there were no differences in spatial bias in the difference score, and no differences in search patterns across the place, cue and “other” quadrants (Fig. 2C-D). Following testing brains were immunostained for the immature neuronal marker doublecortin and qualitative analysis of 2-3 sections from each mouse revealed immature DCX+ neurons throughout the dentate gyrus of all WT mice but extensive loss of adult-born neurons in all of the TK+ mice. Thus, merely inhibiting adult neurogenesis does not influence hippocampal search strategies in the cue-place strategy bias test.

### Running promotes spatial search strategies independently of adult neurogenesis

To test whether a pro-neurogenic factor, exercise, alters search strategies we gave WT mice continuous access to running wheels at 8 weeks of age for 4 weeks. Control mice were not given access to running wheels and both groups were then tested on the cued water maze paradigm (N = 8-9/group). Running did not alter latency to reach the platform in the acquisition phase (Fig. 3A) and did not alter swim speeds (controls 19.9 cm/s, runners 19.6 cm/s; ANOVA effect of running F_1,15_ = 0.3, P = 0.6). However, running did increase spatial search on the probe trial. Running modestly increased the time spent searching in the place quadrant and decreased time spent searching in the cued quadrant, leading to a significant increase in spatial bias (Fig. 3B-C). Specifically, unlike controls, runners spent significantly more time searching in the place quadrant than the cued quadrant. Runners also spent similar amounts of time in the cued and “other” quadrants. In a separate experiment we tested control and running mice on the cued water maze followed by a spatial probe trial where the cue was absent (Fig. 3D-F; N = 10-12/group). Here, running did not enhance spatial search, indicating that the effects of running are restricted to situations where place and cue strategies lead to competing behavioral responses. Notably, unlike all other experiments, mice spent less time searching in the “cued” quadrant (SE) than the “other” quadrants, likely because the cued/SE quadrant is located furthest from the place quadrant and in the absence of a cue there is no motivation to explore this region of the pool.

To test whether running promotes a spatial search bias by enhancing adult neurogenesis, we gave WT and TK mice access to a running wheel or let them remain sedentary in their home cage (N = 9-12/group). Mice were injected with BrdU 8 days after running wheels were introduced. After 4 weeks of running or sedentary conditions, wheels were removed from the runners’ cages and mice were tested in the water maze the following day. Neither running nor inhibition of neurogenesis altered acquisition performance in the cued water maze (Fig. 4A-B) or swim speeds (WT control 20.1 cm/s, WT runner 19.4 cm/s, ANOVA effect of running F_1,16_ = 1.2, P = 0.3; TK control 20.6 cm/s, TK runner 20.0 cm/s; ANOVA effect of running F_1,20_ = 1.7, P = 0.2). In the probe trial, quadrant search patterns revealed that running increased spatial search bias in WT mice as previously observed. However, running also promoted spatial search strategies in TK mice. In both genotypes, running modestly increased time spent in the spatial quadrant and decreased time spent in the cued quadrant (Fig. 4C). Both WT and TK runners also displayed increased spatial bias in the quadrant difference score, with no difference between genotypes (Fig. 4D). Mice were perfused the following day and quantification of BrdU+ cells (now 30 days old) revealed that running significantly increased neurogenesis in WT mice but not in TK mice (Fig. 4E-F). Thus, running promotes spatial search strategies when both spatial and cued responses offer possible solutions, but this enhanced spatial bias is not dependent upon elevations of hippocampal neurogenesis.

**Figure 4:**
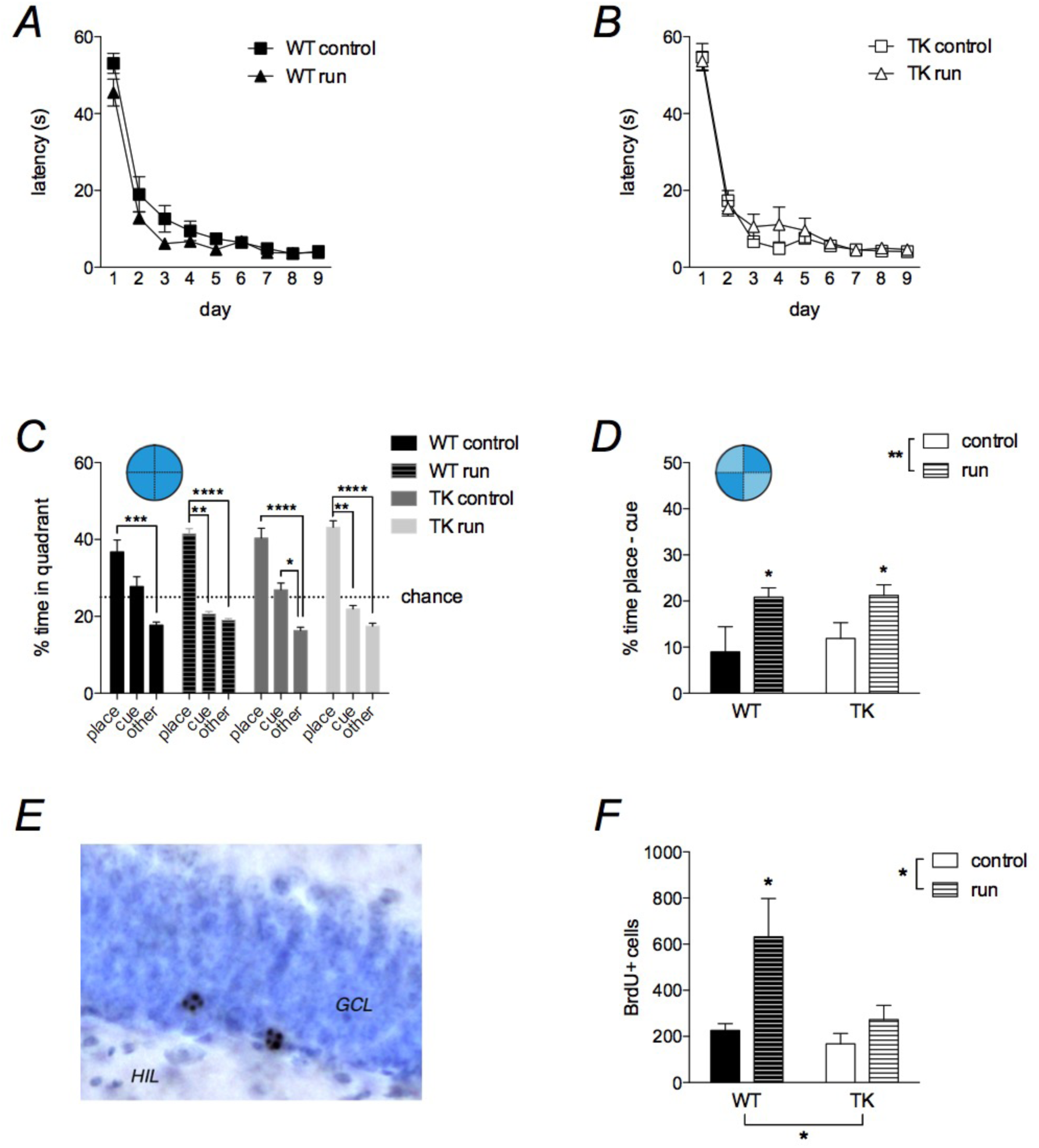
Running promotes spatial bias independently of adult neurogenesis. <u>**A**</u>) Running nonsignificantly reduced escape latencies in WT mice (ANOVA F_1,16_ = 3.9, P = 0.06). <u>**B**</u>) Running did not alter acquisition performance in TK mice (ANOVA, F_1,20_ = 0.5, P = 0.5). <u>**C**</u>) Probe trial quadrant search patterns. Running tended to bias search towards the place quadrant and away from the cued quadrant (Kruskal Wallis tests P < 0.0001; Dunn’s multiple comparisons tests *P < 0.05, **P < 0.01, ***P < 0.001, ****P < 0.0001). There was a trend for greater search time in the cued quadrant than the “other” quadrants in WT control and TK run groups (0.10 < P < 0.05). <u>**D**</u>) Probe trial quadrant difference scores. Running promoted spatial search strategies in the cue-place probe test, in both WT and neurogenesis-deficient TK mice (effect of running F_1,36_ = 9.7, **P < 0.01; effect of genotype F_1,36_ = 0.2, P = 0.6. There was no genotype x running interaction (F^1,36^ = 0.14, P = 0.7) and independent t-tests confirmed that running groups from both genotypes displayed greater spatial bias than their respective controls: WT T_16_ = 2.1, *P < 0.05, TK T_20_ = 2.3, *P < 0.05). <u>**E**</u>) BrdU+ cells in the dentate gyrus of an intact, WT, mouse. HIL, hilus; GCL, granule cell layer. <u>**F**</u>) Running increased BrdU+ cell number in WT but not TK mice (effect of running F_1,35_ = 8.1, **P < 0.01, effect of genotype F_1,35_ = 5.5, *P < 0.05; WT runners vs all other groups *P < 0.05, TK controls vs TK runners P = 0.8). N = 9-12/group.

## DISCUSSION

Here we used a dual solution water maze task to investigate how adult neurogenesis and exercise impact place vs. response learning. While the exact function of adult-born dentate gyrus neurons remains to be determined, they may broadly support hippocampal functions in cognition and spatial processing. We therefore first hypothesized that reducing neurogenesis would compromise hippocampal-dependent place learning and increase reliance on striatal-dependent cued search strategies. Contrary to our predictions we found that ablating neurogenesis did not alter search strategy bias in the cue-place probe test. Second, we expected that increasing neurogenesis with exercise would promote hippocampal function and the adoption of spatial search strategies.

Running, a potent neurogenic stimulus, did promote hippocampal-dependent spatial strategies but this occurred even when neurogenesis was inhibited. Thus, while neurogenesis can contribute to the cognitive effects of exercise (Clark et al., 2008), our current findings as well as previous work (Meshi et al., 2006) indicate that neurogenesis is not always required, and suggest that such conclusions should be tempered in the absence of direct tests.

In our paradigm, spatial bias increased with additional days of training. This would appear to be at odds with previous work showing that hippocampal place strategies dominate initially but dorsal striatal response strategies dominate with additional training (Packard and McGaugh, 1996; Chang and Gold, 2003). These previous studies compared place and response strategies in a plus maze competition test, where rats are released from a constant location and trained to navigate to a food reward that is also in a constant location. During the competition test rats were released from the opposite location of the maze to evaluate place vs response strategies. Compared to our water maze paradigm, plus maze trajectories occur along fixed paths with few options (left or right). The navigational simplicity of the plus maze may therefore favor place strategies relatively early in training, even before the spatial environment is fully learned. In contrast, in our water maze paradigm mice were likely still acquiring relevant spatial (and possibly cue and procedural) knowledge over the 9 days of training. This is suggested by trends in quadrant and zone preferences: with training mice spent less time searching in the “other” areas of the pool that never contained the platform or the cue. Also, while time spent in the cued quadrant decreased with additional days of training, time spent in the cued 20cm zone tended to increase with days of training, indicating a refinement in cued search behavior. Thus, we suspect additional training is required for the switch from place to cued search strategies in our water maze paradigm.

### Identifying spatial memory functions of adult neurogenesis

The precise role of adult neurogenesis in spatial learning remains uncertain. A number of studies have found that blocking adult neurogenesis leads to deficits in spatial water maze learning and memory (Snyder et al., 2005; Imayoshi et al., 2008; Deng et al., 2009; Lemaire et al., 2012) whereas others have found no impairments (Shors et al., 2002; Madsen et al., 2003; Saxe et al., 2006; Jaholkowski et al., 2009; Groves et al., 2013). While the exact reason for these discrepancies remains unclear, a number of differences between studies provides some clues as to the function of neurogenesis in spatial memory. For example, studies that have examined long-term retention of spatial water maze memory have found deficits that were not observed during initial acquisition (Snyder et al., 2005; Ben Abdallah et al., 2013), but see (Saxe et al., 2006). Spatial memory functions may also be dependent on the age of the neurons that are ablated, with a study in mice implicating a role for immature neurons (Deng et al., 2009) and a study in rats implicating more mature adult-born neurons (Lemaire et al., 2012). Other studies have also noted species differences in neurogenesis which could impact behavioral functions (Snyder et al., 2009a; Trinchero et al., 2015). Spared memory in neurogenesis-deficient animals could be due to compensation from other neurons in the dentate gyrus, hippocampus or elsewhere. Indeed, ablating/silencing new neurons impairs memory when the disruption occurs after learning but not before (Arruda-Carvalho et al., 2011; Gu et al., 2012). Only when new neurons are functional at the time of learning are they recruited into the memory trace (Stone et al., 2011). Presumably, when they are absent at the time of learning, other dentate gyrus neurons may become engaged, as has been recently shown in a study of acute silencing and compensatory neuronal recruitment (Stefanelli et al., 2016).

The current study tested a related question: can removing immature neurons shift compensatory functions to the dorsal striatum? Immediate-early gene studies suggest activity in the hippocampus vs. dorsal striatum is not fixed, but may shift depending on the individual or the optimal strategy for solving a task (Colombo et al., 2003; Gill et al., 2007). Furthermore, neurogenesis-deficient mice show less spatially-specific patterns of navigation, suggesting distinct strategy use (Garthe et al., 2009; 2014). Thus, by providing a visible cue it is conceivable that dorsolateral striatum neurons, rather than hippocampal neurons, might compensate for a lack of hippocampal neurogenesis. This would weaken hippocampal encoding during training, and reduce spatial bias on the cue-place probe test. However, we found that blocking neurogenesis did not alter probe trial search strategies, suggesting that older/other hippocampus neurons supported learning of the spatial platform location even though it was cued, and did not require the hippocampus to locate. This finding is consistent with evidence that hippocampal neurons process spatial information similarly regardless of whether a hippocampal or striatal strategy is used (Mizumori et al., 2004; Yeshenko et al., 2004), as well as findings that the hippocampus encodes contextual memories during striatum-dependent learning tasks (McDonald et al., 2002). To rule out a role for neurogenesis in place vs. cue bias future experiments may consider using weaker training paradigms that might depend on neurogenesis to a greater extent (Drew et al., 2010). Also, post-training manipulations of new neurons could be employed to prevent potential compensation from pre-existing hippocampal neurons during the learning process.

### Exercise and spatial bias

In humans, structural and functional differences in the hippocampus and striatum are associated with specific navigational strategies. London taxi drivers, who flexibly navigate a complex spatial environment, have greater posterior hippocampal volumes compared to bus drivers, who follow fixed routes (Maguire et al., 2006). Training on place vs. response variants of the water maze causes volume increases in the hippocampus vs. striatum, respectively (Lerch et al., 2011). In dual solution navigational paradigms, hippocampal activity and volume correlates with spatial strategy use and striatal activity and volume correlates with response strategy use (Iaria et al., 2003; Bohbot et al., 2007). The running-induced spatial bias may therefore be due to non-neurogenic enhancements of hippocampal function. Indeed, other studies have also reported that running and/or enriched environment can improve spatial memory (Meshi et al., 2006) and reduce anxiety-related behaviors (Schoenfeld et al., 2016) even when adult neurogenesis is absent. In rodents, exercise increases hippocampal spine density and dendritic complexity (Redila and Christie, 2006; Stranahan et al., 2007), blood flow (Pereira et al., 2007) and synaptic plasticity (van Praag et al., 1999a). Less is known about exercise effects on striatal function but one report indicates that exercise enhances both place and response learning in single solution tasks (Korol et al., 2013). However, in adult humans, exercise increases hippocampal but not caudate/dorsal striatum volumes (Erickson et al., 2011) suggesting greater effects on hippocampal processing, consistent with our findings. Our observation that running did not enhance spatial preference when the cue was absent would seem to argue against a spatial enhancement effect. However, a more likely explanation is that enhanced place memory could not be detected when a spatial strategy offers the only effective solution. Indeed, control mice performed very well in the uncued probe trial. We propose that weaker spatial representations in controls, relative to running mice, only became apparent once there was a conflict between cued and place responses.

An alternative explanation is that exercise promoted spatial bias by altering interactions between brain regions. For example, the prefrontal cortex coordinates inputs from both the hippocampus and dorsal striatum to decide optimal behavior (Mizumori and Jo, 2013; Dahmani and Bohbot, 2015).

Enhancing prefrontal function with exercise (Prakash et al., 2011; Brockett et al., 2015) may support the use of cognitive, less automated, strategies. Additionally, the amygdala promotes response strategies, particularly under stress (Packard and Gabriele, 2009; Leong and Packard, 2014; Leong et al., 2015). Exercise may therefore promote spatial bias by reducing stress-related amygdala modulation of hippocampus and dorsolateral striatum function. Changes in neuromodulator signalling could also promote hippocampal strategy choice. Region-specific differences in cholinergic and dopaminergic signalling contributes to hippocampal vs. striatal strategies in dual solution tasks (Packard and White, 1991; McIntyre et al., 2003). Reduced glucocorticoid-norepinephrine signalling (Goodman et al., 2015) or enhanced estrogen signalling (Korol and Pisani, 2015) are also candidates for exercise-mediated spatial bias. Finally, animals with hippocampal damage can demonstrate a limited degree of spatial learning (Eichenbaum et al., 1990; Whishaw et al., 1995; Day et al., 1999). Thus, it is possible that exercise may enhance spatial processing in non-hippocampal regions, such as the neocortex (Teixeira et al., 2006; Tse et al., 2011; Czajkowski et al., 2014) or dorsal striatum (Yeshenko et al., 2004; Ferbinteanu, 2016), both of which have been shown to contribute to spatial memory under certain conditions.

### Conclusions and relevance for psychiatric disorders

To our knowledge, this is the first study to examine whether exercise influences spatial vs. response strategies when subjects are given a choice in a dual solution task. Our findings are relevant for a number of disorders that are characterized by alterations in habit behavior. For example, it has been proposed that the dorsal striatum may enhance responding to threatening cues in posttraumatic stress disorder, while hippocampal dysfunction impairs memory for the details of traumatic experiences, leading to fear and anxiety responses in inappropriate contexts (Goodman et al., 2012). Imbalances between the hippocampus and striatum may also bias individuals with addiction, obsessive compulsive disorder and Tourette’s Syndrome towards striatal dependent habit behaviors (Bohbot et al., 2013; Goodman et al., 2014). Our findings suggest that exercise could be beneficial by biasing towards hippocampal-dependent cognitive behaviors, thereby promoting behavioral flexibility and more healthy decisions.

